# Metformin regulates bone marrow stromal cells to accelerate bone healing in diabetic mice

**DOI:** 10.1101/2023.04.05.535802

**Authors:** Yuqi Guo, Jianlu Wei, Chuanju Liu, Xin Li, Wenbo Yan

## Abstract

Diabetes mellitus is a group of chronic diseases characterized by high blood glucose levels. Diabetic patients have a higher risk of sustaining osteoporotic fractures than non-diabetic people. The fracture healing is usually impaired in diabetics and our understanding of the detrimental effects of hyperglycemia on fracture healing is still inadequate. Metformin is the first-line medicine for type-2 diabetes (T2D). However, its effects on bone in T2D patients remain to be studied. To assess the impacts of metformin on fracture healing, we compared the healing process of closed wound fixed fracture, non-fixed radial fracture, and femoral drill-hole injury models in T2D mice with and without metformin treatment. Our results demonstrated that metformin rescued the delayed bone healing and remolding in T2D mice in all the three injury models. The proliferation, osteogenesis, chondrogenesis of the bone marrow stromal cells (BMSCs) derived from WT and T2D mice treated with or without metformin were compared *in vitro* and *in vivo* by assessing the subcutaneous ossicle formation of the BMSC implants in recipient T2D mice. *In vivo* treatment with metformin to T2D mice could effectively rescue the impaired differentiation potential and detrimental lineage commitment of BMSCs, caused by the hyperglycemic conditions. Moreover, the Safranin O staining of cartilage formation in the endochondral ossification under hyperglycemic condition significantly increased at day 14 post-fracture in T2D mice receiving metformin treatment. The chondrocyte transcript factor SOX9 and PGC1α, important to maintain chondrocyte homeostasis, were both significantly upregulated in callus tissue isolated at the fracture site of metformin-treated MKR mice on day 12 post-fracture. Metformin also rescued the chondrocyte disc formation of BMSCs isolated from T2D mice. Taken together, our study demonstrated that metformin facilitated bone healing, bone formation and chondrogenesis in T2D mouse models.

## Introduction

Diabetes mellitus is a group of chronic diseases characterized by high blood glucose levels. It is estimated that more than 347 million people worldwide currently have diabetes and many more people are estimated to become diabetic soon [1]. Among the diabetic patients, more than 90% are suffering from Type-2 diabetes (T2D), which is caused by insulin resistance in peripheral tissues. Many tissues including the skeleton will be adversely affected by hyperglycemia if not controlled [2]. Both Type 1 diabetes and T2D are associated with an increased risk of osteoporosis and fragility fractures [2]. It is recognized that oral antidiabetic medicines affect bone metabolism and turnover [2]. As an insulin sensitizer, patients with T2D are frequently prescribed with metformin. In T2D patients, metformin treatment was associated with a decreased risk of bone fracture [3]. The osteogenic effects of metformin have been documented in both cellular and rodent models. Metformin promoted osteoblast differentiation and inhibited adipocyte differentiation in rat bone marrow mesenchymal stem cells culture [4]. In rat primary osteoblasts culture, metformin increased trabecular bone nodule formation [5]. In ovariectomized rats, metformin was also shown to improve the compromised bone mass and quality [6]. Furthermore, in streptozotocin-induced Type-1 DM model, metformin stimulated bone lesion regeneration in rats [7]. However, to date, there is no study on the skeletal effects of metformin in a T2D model. A recent study found that the incidence of total knee replacement over four years was 19% lower among patients with type 2 diabetes who were regular metformin users, compared with non-users [8]. In addition to reducing glucose levels, metformin may modulate inflammatory and metabolic factors, leading to reduced inflammation and plasma lipid levels [8]. Better knowledge of how metformin treatment influences skeletal tissues under T2D condition is of great clinical relevance in view of the fast-growing population of patients with T2D.

MKR mouse model was generated and characterized by LeRoith and colleagues and expresses a dominant negative mutant of human IGFI receptor specifically in skeletal muscles [9]. The expression of this dominant negative mutant human IGFI receptor decreased glucose uptake and causes insulin resistance, the MKR transgenic mouse rapidly develops severe diabetes [9]. This mouse model has been widely used in T2D research. In this study, the effects of metformin on bone cell lineage determination and regeneration was characterized using MKR mouse model.

## Materials and Methods Animals

### Bone injury models

#### 1. Femoral closed fracture model

The gravity-induced Bonnarens and Einhorn fracture model was adapted here as previously described in order to establish a standard closed fracture [10, 11]. Briefly, 12- week old WT and MKR male mice were anesthetized with a Ketamine/ Xylazine cocktail, and a 1 cm sagittal incision was made at the right knee beneath the patella. A 3/8-inch length 27-gauge needle was inserted into bone canal right between medial and lateral condyles. The needle end was cut, and the blunt end was pushed forward and buried between medial and lateral condyles to avoid tissue damage afterwards. With the fixative needle inside of the femur canal, animal was moved to the fracture apparatus. The right femur was placed over the two supports, and the blunt guillotine blade was dropped from a pre-tested height onto the femur, to created sufficient force to cause the fracture. The wound was then closed with suture, and mice were randomly assigned into vehicle (PBS) or metformin (Met, 200mg/kg BW) daily treatment groups.

#### 1. 2. Radius non-union fracture model

Male WT and MKR mice of 12-week-old were anesthetized using a Ketamine/ Xylazine cocktail. A 0.5 cm coronal incision was made over the right radius. The brachioradialis and pronator teres were carefully separated with blunt surgical instruments to reveal the radius. A super sharp Stevens Tenotomy Scissor was used to cut at the middle of the radius and created the non-union radial fracture. The wound was then closed, and mice were randomly assigned into PBS or metformin daily treatment groups.

#### 1. 3. Femoral drill hole model

WT and MKR male mice of 12-week-old were anesthetized using a Ketamine/ Xylazine cocktail. A 1 cm coronal incision was made over the right lateral femur. Quadriceps were carefully separated with blunt surgical instruments to reveal the femur. A drill bit (#66) was used to create a 0.8 mm diameter hole on the femur. After closing the wound, mice were randomly assigned into PBS or metformin daily treatment groups.

### Tissue Collection and Processing

The micro computed tomography (μCT) and bone histomorphometry were utilized to assess static and dynamic indices of bone structure and formation. Briefly, bone injuries were introduced in animals as described above, PBS and metformin treatment was administrated daily for indicated time. After sacrificing the animal, injured bone samples were fixed in 10% buffered formalin for 48 hours, then rinsed with PBS before being analyzed by μCT. Bones were evaluated using a SkyScan 1172 high-resolution scanner (Brucker, Billerica, MA, USA) with 60 kV voltage and 167μA current at a 9.7 μm resolution and reconstructed using NRecon (V.1.6.10.2.). Whole scanned region was included as VOI (volume of interest) [12] from femoral fracture model to generate a general 3D view of the femur fracture cite. In the radius fracture model, a total of 3 mm (311 transverse anatomic slides) radius including the entire injury site was selected as VOI. In order to analyze only the fracture callus bone parameters, two sets of regions of interest [9] were manually drawn within each VOI slides. The first set traced the external fracture callus (inclusion ROI) and the second set traced the cortical bones within the callus (exclusion ROI). In the femur drill hole model, a round shaped ROI with 0.611 mm diameter (63 pixels) was surrounded in the center of injury hole throughout the depth where callus exhibit to generate the VOI consist of callus within the drill cite. VOI from each animals were analyzed with CTan (V.1.13.2.1.) to calculate the following morphometric parameters: bone mineral density (BMD), relative bone volume (BV/TV), trabecular thickness (Tb.Th), trabecular separation (Tb. Sp), porosity, and total pore space. CTVox (V.3.3.0.0.) was used to generate the 3 spatial images of the VOI.

Callus bridging score was ranking from 0 to 3. (0: no bridging; 1: less than 33% bridging; 2: more than 34% but less than 66% bridging; 3: more than 67% till complete bridging). Callus bridging ratio was calculated by each animals’ callus bridging score/3. For double labeling experiment, ten-week-old male WT and MKR mice were randomly assigned to receive PBS or metformin daily injections for 14 days. Mice also received intraperitoneal injections of calcein (10 mg/kg) on the 5^th^ day and alizarin red (15 mg/kg) on the 12^th^ days during the 14-day period. For histomorphometry and peripheral quantitative computed tomography, femurs were preserved in 70% ethanol until they were processed for plastic embedding in methyl methacrylate resin or decalcified for paraffin embedding.

Blood was collected by cardiac puncture after euthanasia, left at room temperature for 30 min before centrifuging at 200 g for 10 min to separate sera.

### Cell culture and analysis

As previously described, after 14-day treatment of PBS or metformin *in-vivo*, WT and MKR mice were sacrificed and proceeded with bone marrow primary cell culture. Excessive tissue was removed from femur and tibia from each mouse, and quickly rinsed with 70% ethanol and then followed by triple cold PBS wash to ensure sterility. Bone marrow cells were flushed out using cold PBS and broken down into individual cell suspension by passing through a 19-gauge needle for 5 times. Cells were then cultured in MEM Alpha Modification (α-MEM) medium supplemented with 10% Fetal Bovine Serum (R&D Systems, USA), 100 μg/ml streptomycin, and 100 units/ml penicillin (Gibco, Grand Island, NY, USA) in a 37 °C, 5% (v/v) CO2, humidified incubator. After removed non-adherent cells, the adherent cells were cultured as bone marrow stromal cells (BMSCs) for 7 days and sub-cultured for following assays:

#### 1. Colony-forming unit fibroblastic assay (CFU-F)

Isolated BMSCs were seeded at 100, 500, and 1000 cells/well in 6-well plate, followed by 10 days culture with α-MEM complete medium. Cells were fixed with 10% buffered formalin, washed 3 times with PBS followed by staining with 0.25% wt/vol crystal violet solution. Plate images were captured using ChemiDoc XRS System (BioRad Laboratories, Inc. Hercules, CA, USA) and analyzed with ImageJ.

#### 1. 2. Osteoblast differentiation

Isolated BMSCs were seeded at 6.4 x 10^4^ cells/well into 6-well plates, and cultured with osteogenic medium containing α-MEM complete medium supplement with 50 µg/mL L-Ascorbic acid-2-phosphate and 10mM β-glycerophosphate for 2∼3 weeks followed by ALP and von Kossa staining. Briefly, after 2 weeks of osteogenic differentiation, cells were fixed and checked for alkaline phosphatase activity using ALP kit (86R-1KT, Sigma) following manufacture’s protocol. After 3 weeks under osteogenic culture, cells were fixed with 95% ethanol and rehydrated through gradient ethanol to water. Cells were then carefully washed with water after being incubated with 5% silver nitrate solution at 37 °C for one hour, exposed under UV light for 10 minutes. All plates’ images were captured using ChemiDoc XRS System, and ALP positive area and mineralized regions were measured by ImageJ software.

#### 1. 3. Chondrocytes differentiation and disc formation

Isolated BMSCs were further cultured and passaged twice using standard growth media of DMEM+10% FBS in order to enrich the cell number. A cell solution of 1.6x10^7^ cells/mL was generated using StemPro™ Chondrogenesis Differentiation Kit (ThermoFisher), and 5 µL droplets were applied in the center of 48-well plate wells for micro-mass culture. 3 hours later, warmed chondrogenic medium were overlayed over the micro-mass and the formation of osteogenic pellets were observed after 3 days of culture.

### *In vivo* Ossicle formation assay

As described above, BMSCs from different treatment groups were obtained and cultured using standard growth media of DMEM+10% FBS with customized glucose level. We measured and calculated mean value of recipient MKR mice blood glucose level (N=4, average glucose level = 436 mg/dL) and prepared the ex-vivo culture medium accordingly to avoid glucose level change from ex-vivo culture to the body fluid of the recipient MKR mice. Briefly, BMSCs from PBS or metformin treated mice were growing to 90% confluent, then 1.0x10^8^ cells/mL cells solution was obtained and 20 µL of this cell solution was soaked in a 4mm x 4mm gel foam and grafted at the flank of the recipient MKR mice subcutaneously. Four weeks after the implantation, the gel foams were dissected and processed for histological assays.

### Histology

Femurs from fracture models and bone ossicles were decalcified using 10% EDTA for 2 weeks. EDTA solution was refreshed every other day for the best decalcification efficacy. Tissues were then processed through automatic tissue processor, followed by paraffin embedding. H&E, safranin O, and Mason’s Trichrome staining were performed respectively.

### Statistics

We used ANOVA analysis when the study subjects included more than two groups, followed by the Bonferroni t-test. We used the two-tailed Student’s t-test to compare the difference between two experimental groups. A value of P<0.05 was considered to be statistically significant. Bars in figures represent the mean ± SEM unless stated otherwise.

## Results

### Metformin promotes healing in fracture models under hyperglycemic condition

In order to evaluate the fracture healing which involves endochondral ossification in mice, we adapted this well accepted Bonnarens and Einhorn fracture mouse model (Fig 1A). Animals were sacrificed on day 14, 23, or 31 post-fracture representing the inflammatory stage, the endochondral stage, and the remodeling stage during the femur fracture repair process [13]. As shown in Fig 1B, metformin treated MKR mice exhibited similar healing and remodeling speed to that of the WT groups throughout the entire healing process, and the fractured bones were completely healed on day 31. On the other hand, the healing process in MKR-PBS group was much delayed, with visible amount of callus tissue remained at the fracture site on day 31. Additionally, histological analysis of the callus area (Fig. 1C) disclosed a postponed peak callus formation in MKR-PBS group when compared to WT-PBS, WT-met, and MKR-met groups on day 23 post-fracture, as well as a delayed remodeling on day 31 post-fracture. A quantitative analysis of the callus at the fracture area (Fig. 1D) confirmed that metformin significantly enhanced bone healing in T2D mice comparing to PBS treated MKR mice.

**Figure 1.**
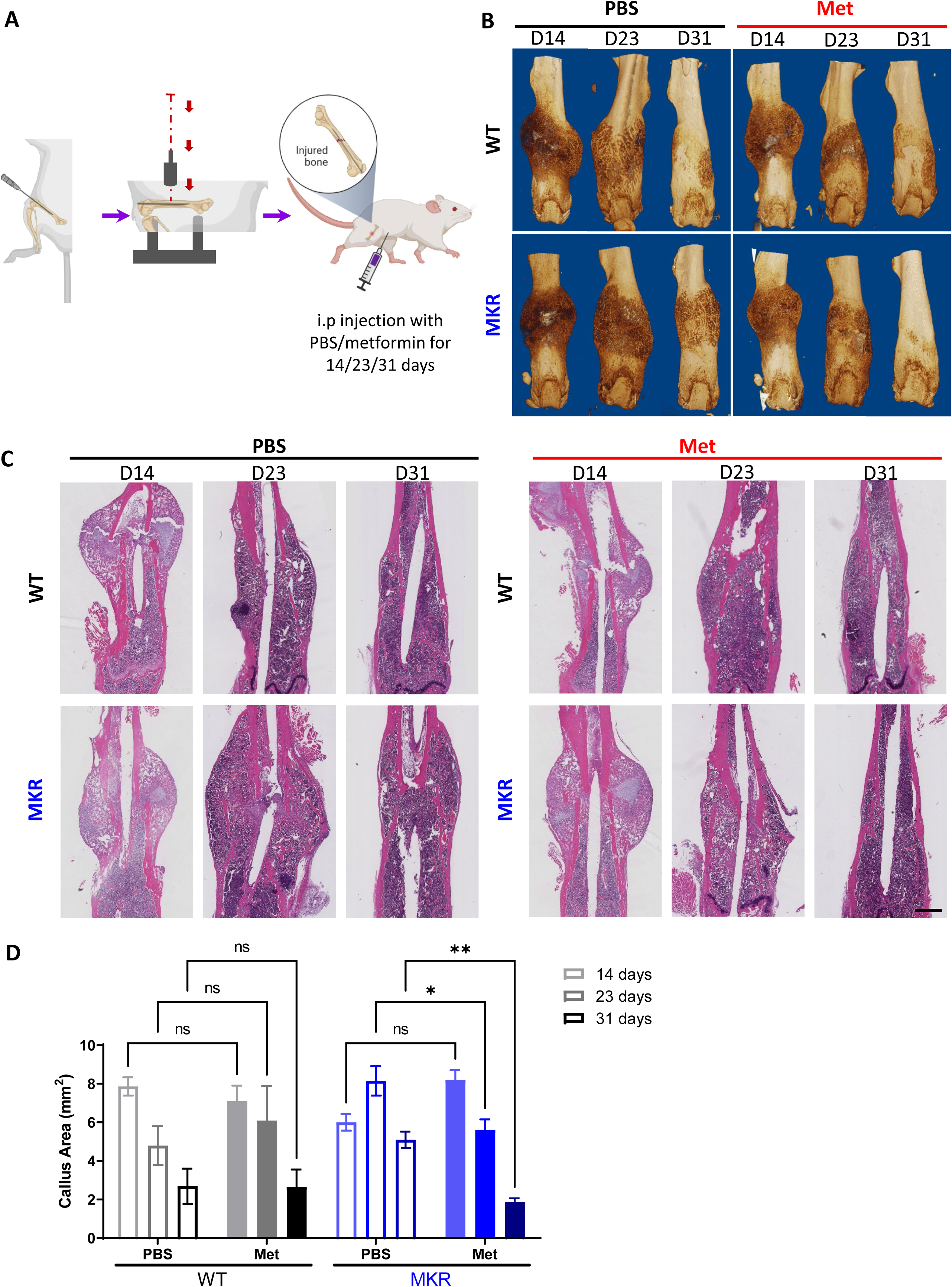
Improved healing in closed transverse fracture. A) Schematic representation of the experimental design. B) Representative µCT images of femurs from each treatment group. C) H&E staining of longitudinal femur sections (scale bar, 1 mm) and D) Fracture callus area analysis from each treatment group. (Bars show mean ± SEM. n= 6∼9, *p<0.05, *** p<0.005).

A similar effect of metformin was also observed in a non-fixed radial fracture model (Fig. 2). After non-fixed radial fracture was introduced, animal was treated with either PBS or metformin for 14 or 23 days (Fig. 2A). In the WT groups at 14 days post-fracture, both PBS and metformin treated animals started to exhibit sufficient amount of callus with the sign of bridging of the fracture ends (Fig. 2B). In the 14-day post-fracture MKR mice, those treated with metformin exhibited more healing callus than the PBS treated ones with significantly greater percentage of callus bridging at the fracture site (Fig. 2C). Bone mineral density was also higher in metformin-treated MKR mice when compared to the PBS-treated MKR mice (Fig. 2D). Bone volume/tissue volume ratio (Fig. 2E) and trabecular thickness (Fig. 2F) were not regulated by metformin in either WT or MKR mice. In the 23 days post-fracture groups, only PBS-treated MKR animals failed to show callus formation and bridging at the fracture site compared to all the other groups (Fig. 2G and 2H), indicating a delayed healing process in MKR PBS group. Bone mineral density levels in metformin-treated MKR mice were similar to the PBS-treated WT mice and were significantly higher than the PBS-treated MKR mice (Fig. 2I). In the 23 days post-fracture groups, the bone volume/tissue volume ratio (Fig. 2J) and trabecular thickness (Fig. 2K) were significantly improved by metformin in MKR mice in comparison to the PBS-treated MKR mice. These data suggest that 23 days of metformin treatment favorably affected bone healing. The beneficial effects of 23 days of metformin treatment on the callus bridging (Fig. 2H), BV/TV% (Fig. 2J), and Tb. Thickness (Fig. 2K) became significant in contrast to the trends of improvement observed in the mice received short term treatment for 14 days (Fig 2C, 2E and 2F).

**Figure 2.**
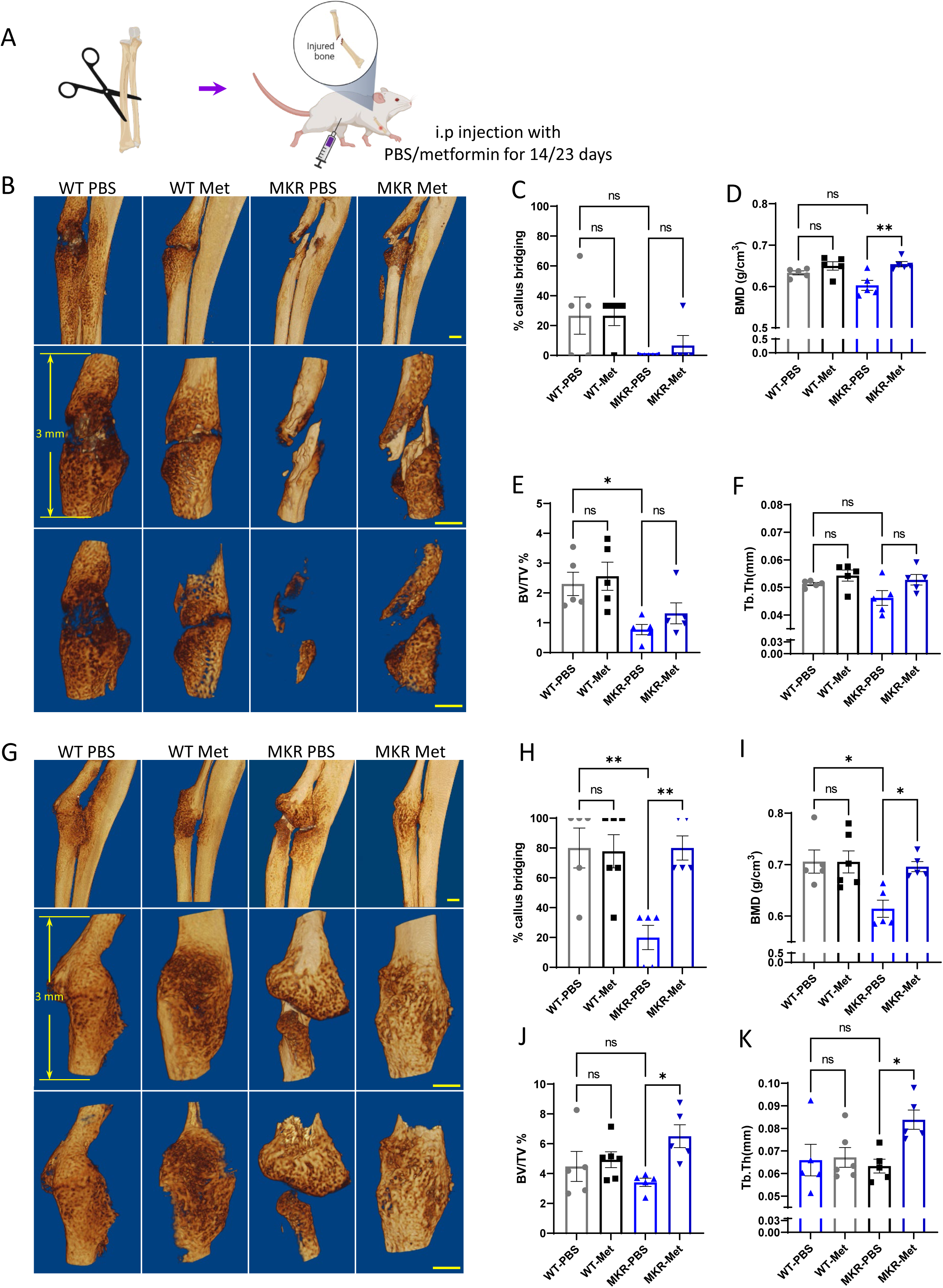
Improved healing in non-fixed radial fracture at two different time points (14 days and 23 days post fracture). A) Schematic representation of the experimental design. B) Representative µCT images of mouse radiuses (top) and fracture sites (bottom) 14 days post-fracture (scale bar, 500 µm). Measured parameters (C-F) by Micro-CT at 14 days post-fracture. C) Percentage of callus bridging (%). D) Bone mineral density (BMD; g/cm^3^). E) Bone volume/tissue volume (BV/TV; %). F) Trabecular thickness (Tb.Th; mm). G) Representative µCT images of mouse radiuses (top) and fracture sites (bottom) 23 days post-fracture (scale bar, 500 µm). Measured parameters (H-K) by Micro-CT at 23 days post-fracture. H) Percentage of callus bridging (%). I) Bone mineral density (BMD; g/cm^3^). J) Bone volume/tissue volume (BV/TV; %) - K) Trabecular thickness (Tb.Th; mm). Results of quantitative µCT data analysis (bars show mean ± SEM.).

### Metformin improves healing in the drill hole injury model under hyperglycemic condition

To further investigate metformin’s effect on bone repair, a drill-hole model was established (Fig. 3A). After 14 days treatment with PBS or metformin, we examined the injury site using µCT scanning. Exterior and interior general view of the femur indicate a non-filled open wound exhibited only in PBS treated MKR group, while all the other three groups showed mainly filled holes indicating a faster healing speed (Fig. 3B-D).

**Figure 3.**
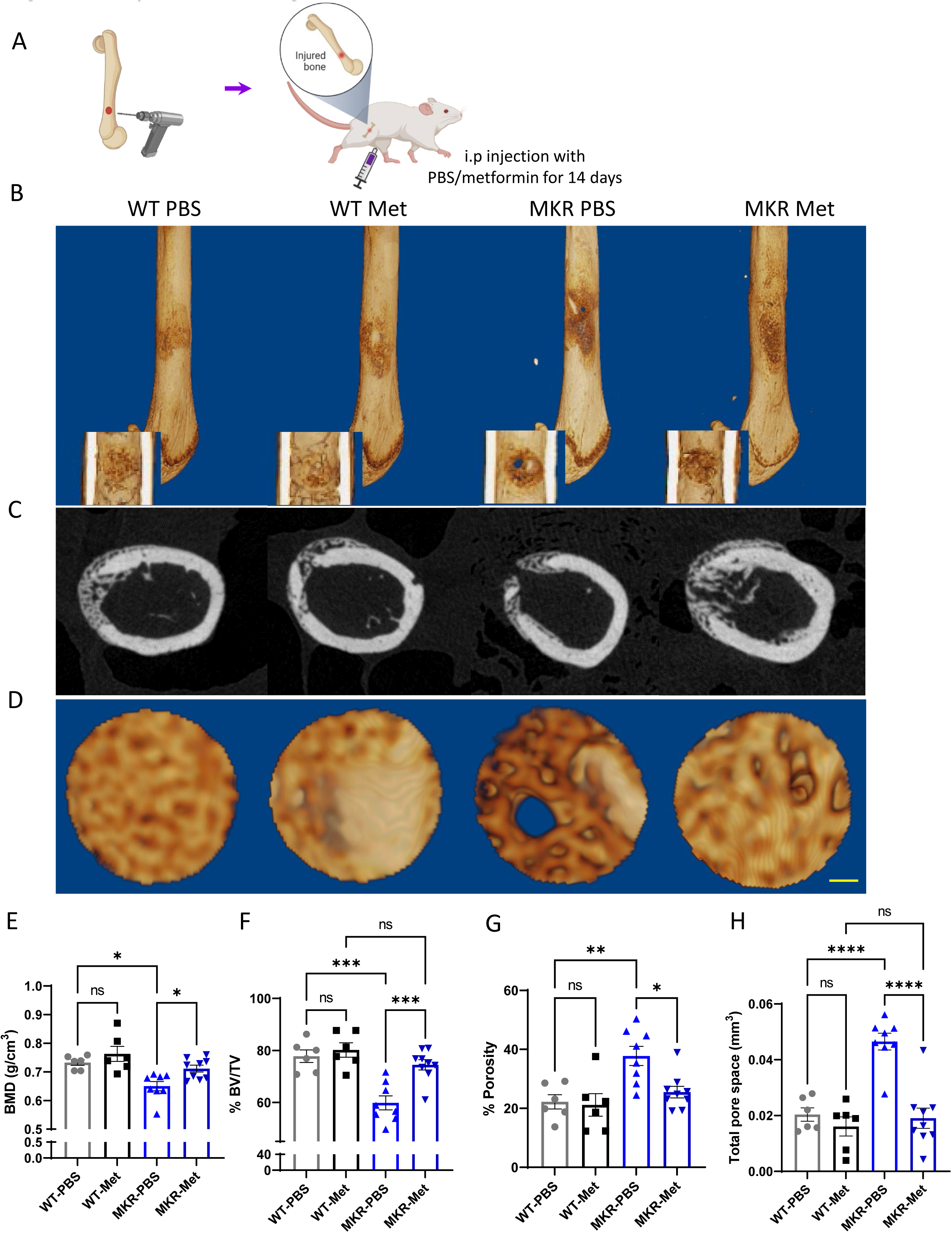
Improved healing in drill-hole bone repair model. A) Schematic representation of the experimental design. B) Representative 3D images of mouse femurs with both exterior and interior general view at 14 days post-surgery. C) Cross plane images at center of the drill site. D) Representative 3D images within the drill site (scale bar, 100 µm) and E)-H) Results of quantitative µCT data analysis (Bars show mean ± SEM. n=6∼9, *p<0.05, **p<0.01, *** p<0.005, **** p<0.0001). E) Bone mineral density (BMD; g/cm3). F) Bone volume/tissue volume ratio (BV/TV; %). G). Porosity (%). H). Total pore space (mm^3^)

Tracing of the drill-hole injury sites presented a detailed view of the callus tissues formed within the drill hole (Fig. 3D). In the WT animals, quantitative analysis of the µCT images showed no difference between the PBS and the metformin treatments. Significant lower BMD (Fig. 3E) and BV/TV ratio (Fig. 3F) were observed in PBS treated MKR mice when compared to the metformin treated ones. The PBS treated MKR mice also demonstrated prominently higher bone porosity (Fig. 3G) and total pore space (Fig. 3H) within the callus tissue, suggesting delayed bone healing and remodeling in MKR mice was rescued by metformin.

### Metformin accelerates bone formation under hyperglycemic condition

In order to examine metformin’s effects on bone formation *in vivo*, we conducted bone Alizarin Red and Calcein Double Labeling injections on WT and MKR mice. Fig. 4A suggested that the florescent labels in WT mice remained the same between PBS and metformin treated animals. In contrast, the distance between the two labels was greater in metformin treated than the PBS treated MKR mice. This observation was supported by serum levels of amino-terminal propeptide of type 1 procollagen (P1NP). P1NP is considered a sensitive marker of bone formation [14] and the ELISA assay was performed using the serum samples collected at 14, 23, and 31 days post femoral fracture. In WT animals, there is no difference in P1NP levels between metformin treated and PBS-treated mice at all three time points (Fig. 4B). On the contrary, we observed significantly higher P1NP levels at all three time points post-fracture in the MKR mice treated with metformin compared those treated with PBS (Fig 4B). Collectively, the data indicate that metformin can promote bone formation only under hyperglycemic conditions.

**Figure 4.**
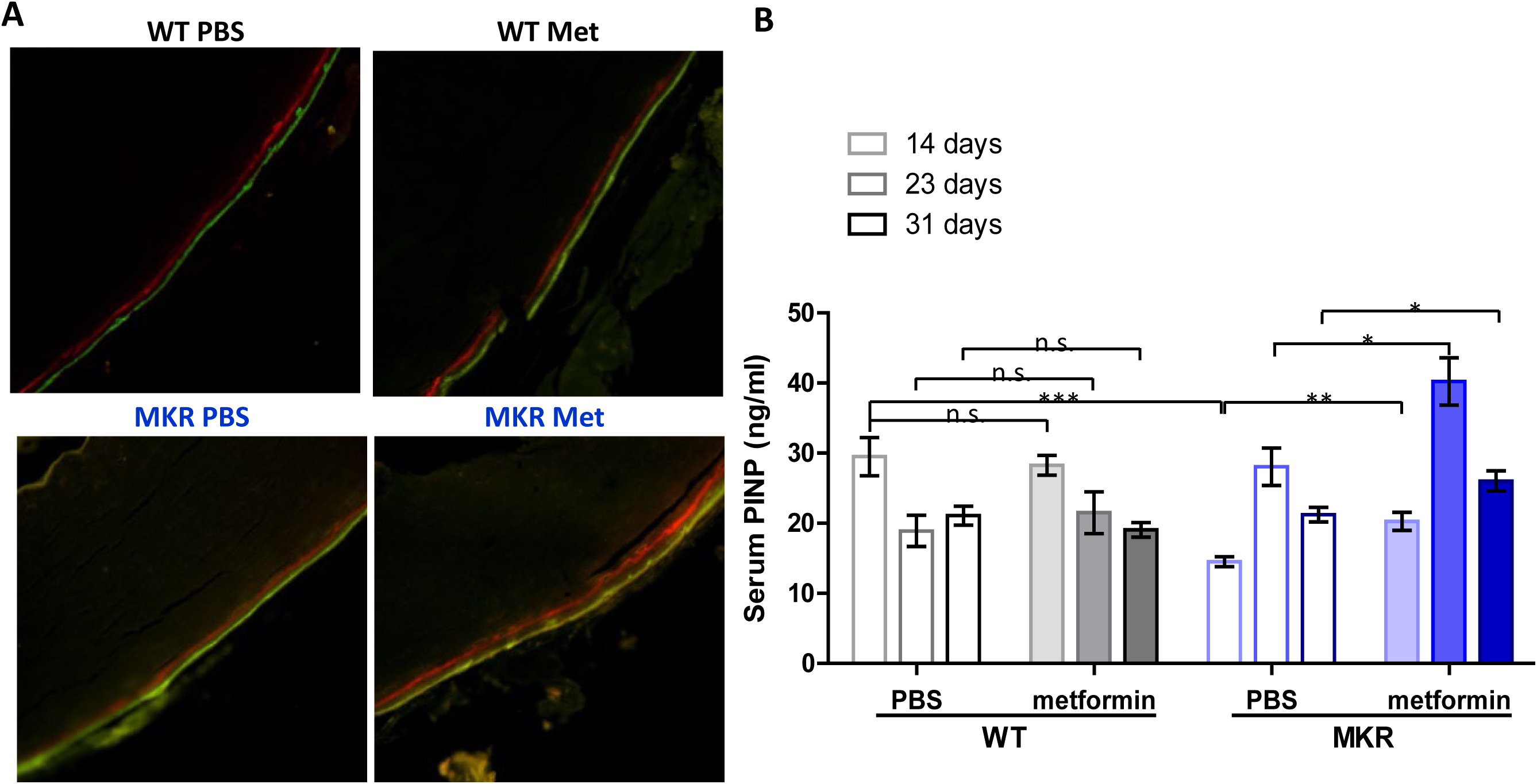
Metformin promotes bone formation. A) Representative images of calcein double-labeling in cortical periosteum of the femur mid-shaft. B) Serum P1NP level (ng/ml) from each treatment group at 14, 23, and 31 days post-femoral fracture. (Bars show mean ± SEM. n=6∼9, *p<0.05, **p<0.01, *** p<0.005)

### Metformin regulates proliferation and lineage commitment of the bone marrow stromal cell (BMSC) in MKR mice

Considering that the multipotent mesenchymal stem/progenitor cells (BMSCs) in bone are critical in maintaining bone quality, function and regeneration, we then tested whether metformin stimulated BMSCs proliferation in MKR mice. As expected, hyperglycemic condition in MKR mice impaired the proliferation of their BMSCs, as indicated by CFU-F staining when compared to WT controls (Fig. 5A). Administration of metformin in MKR mice successfully salvaged the BMSC proliferation capability and brought it back to the equivalent level as observed in WT group. Under all three seeding densities, compromised CFU-F colony formation observed in MKR group can be rescued by metformin treatment as shown in Fig. 5B. However, metformin did not affect the proliferation of BMSCs in WT group when compared to the PBS vehicle. It is noteworthy that metformin was not administrated to the culture. The daily metformin treatment in MKR mice prior to cell isolation appeared to be sufficient to protect the proliferation potential of BMSCs from the detrimental effects of hyperglycemia. BMSCs lineage differentiation potential was also tested by ALP staining and von Kossa staining. After 14 days under osteogenic [1] differentiation, in contrast to the weak ALP activity observed in MKR-PBS group, MKR-met group showed compatible ALP activity to the WT groups (Fig. 5C and 5D). ALP played a critical role in calcium crystallization and mineralization during bone formation, therefor we speculated this aberrant ALP activity of the MKR-PBS group would further lead to an impaired bone mineralization. As expected, bone mineralization was barely detected in MKR-PBS group after 21 days osteoblast differentiation visualized by von Kossa staining and metformin significantly enhanced bone mineralization in MKR mice when compared to the PBS-treated MKR animals (Fig. 5E and 5F). Notably, the BMSCs from metformin treated T2D mice could maintain the improved bone formation feature when re-exposed to the same hyperglycemic levels as in the T2D mice. BMSCs isolated from metformin treated MKR mice and PBS controls were implanted to the recipient MKR mice (Fig. 5G). Masson’s Trichrome staining on the ossicles formed by the BMSCs from metformin treated MKR mice showed greater bone formation than that of PBS treated MKR mice (Fig5 H-I). These results implied that *in vivo* treatment of metformin could effectively rescue the impaired differentiation potential of BMSCs in MKR mice.

**Figure 5.**
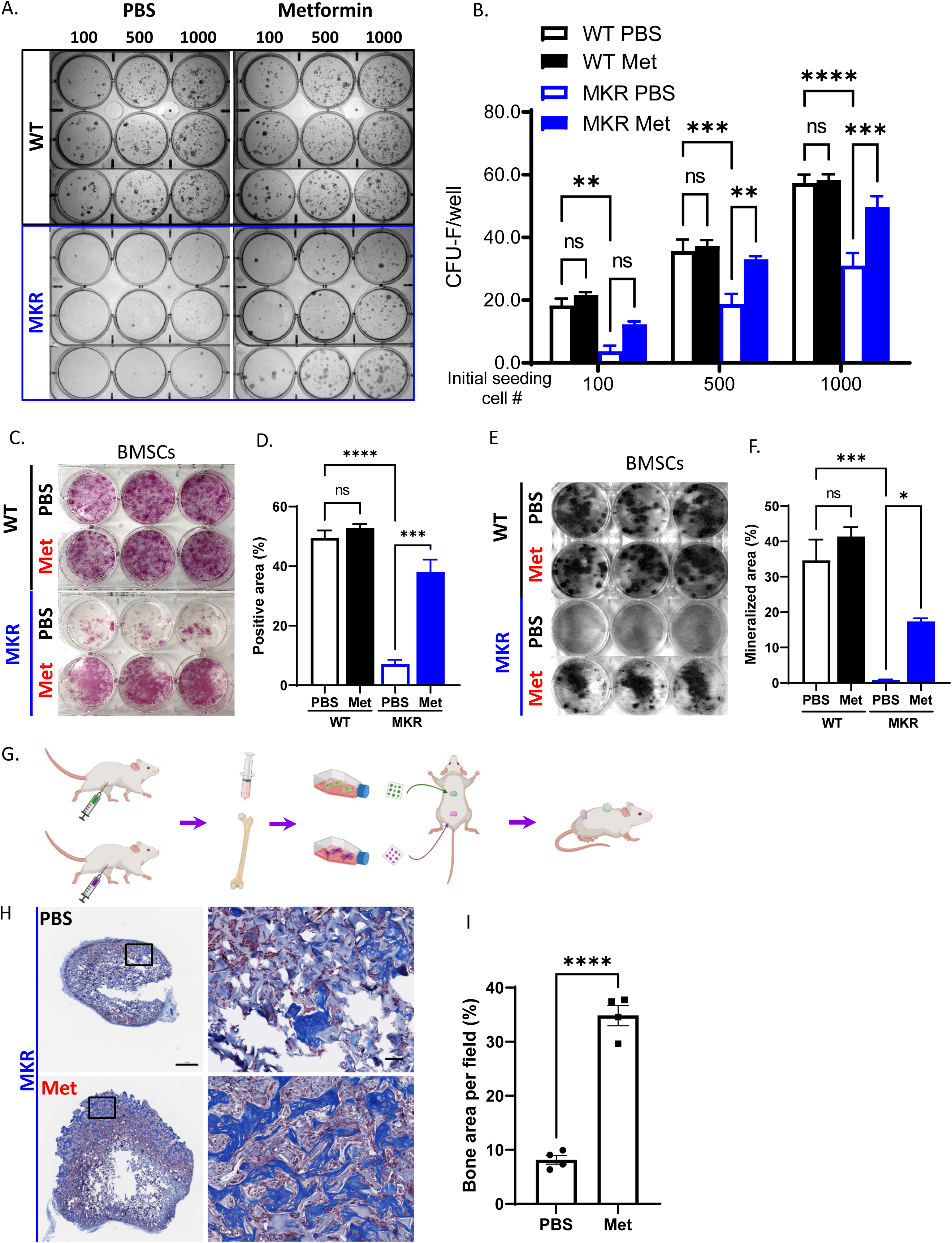
Improved osteogenesis of BMSC from metformin treated MKR mice. A) Primary BMSCs were isolated from animals that were treated with PBS or metformin in-vivo for 14 days and were seeded at indicated density for CFU-F culture. B) Results from quantitative analysis of colony counts measured by ImageJ. C) Primary BMSCs from mice received 14-day treatment of PBS or metformin were plated in 6-well plates and cultured using differentiation medium and were tested for ALP activity, D) total ALP positive area per well were measured by ImageJ. E) von Kossa staining to examine mineralization. F) Calculation of mineralized area. G) Schematic representation of the experimental design. H) Masson’s Trichrome staining on the ossicle sections, and I) percentage of bone area (blue) per field was measured using ImageJ. (Bars show mean ± SEM. n= 6∼9, *p<0.05, **p<0.01, *** p<0.005, **** p<0.0001).

### Improved chondrogenesis of BMSC from metformin treated MKR mice

Chondrogenesis and endochondral ossification are critical steps during the healing process after a bone injury. In order to examine metformin’s effect on chondrogenesis during bone healing, we compared the cartilage deposition within fracture healing sites throughout the healing process (14 days, 23 days, and 31 days post fracture). The cartilage deposition was significantly lower in PBS-treated MKR mice as compared to the PBS-treated WT mice at 14days post fracture (Fig. 6A-B). As expected, no difference between the PBS or metformin treated animals was observed in WT groups. On the other hand, metformin significantly promoted cartilage formation in MKR mice on day 14 post fracture, and the trend continued till day 23 (Fig. 6A-B). By day 31 post fracture, except in the PBS treated MKR mice, no discernible callus remained in any other groups (Fig. 6A). To further investigate if metformin modulates chondrogenesis of BMSCs. BMSCs were isolated for chondrogenesis culture in vitro. Only BMSCs isolated from PBS treated MKR mice failed to form the chondrocyte disc as shown in Fig. 6C after 3 days chondrogenic culture. BMSCs isolated from metformin treated MKR mice could form the chondrocyte disc as well as the WT controls (Fig. 6C). We also harvested the callus tissue at fracture sites on day 12 and day 21 post fracture to examine the expression of genes that contribute to chondrogenesis. At day 12 post fracture, the chondrocyte transcript factor SOX9 was significantly upregulated in metformin treated MKR mice (Fig. S2C). SOX9 is a master transcription factor that play a key role in chondrogenesis[15]. At day 21 post fracture, the PGC1α was significantly upregulated in metformin treated MKR mice (Fig. S2G). PGC1α is required for chondrocyte metabolism and cartilage homeostasis [16, 17].

**Figure 6.**
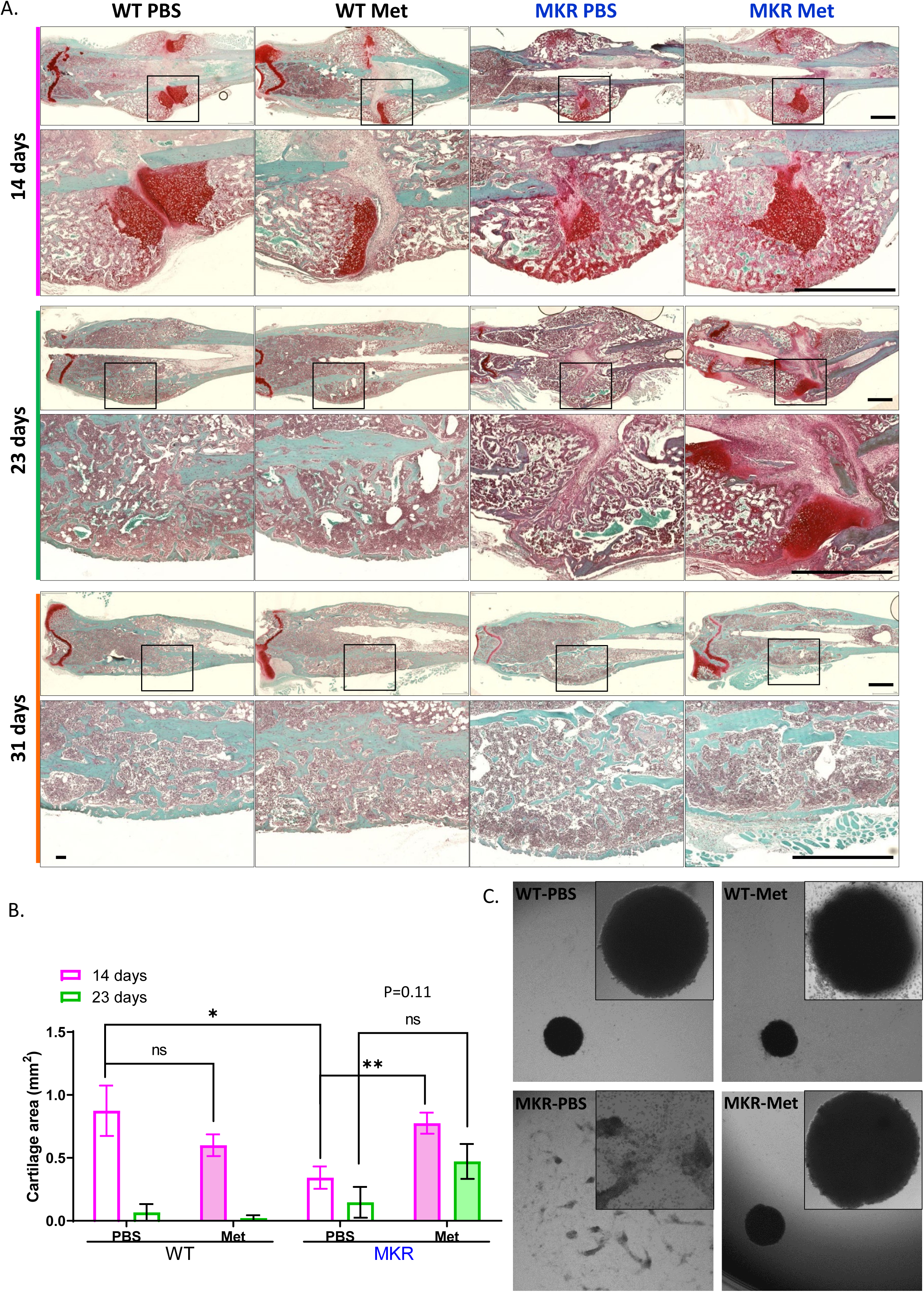
Improved chondrogenesis of BMSC from metformin treated MKR mice. A) Safranin O staining of longitudinal femur sections from femoral fractures at different time points (scale bar, 1 mm). B) Cartilage area [18] at fracture site were measured using ImageJ. C) Chondrocyte pellet culture of BMSCs from PBS or metformin treated animals. Micromass culture were generated by seeding 5 µL of primary BMSCs (1.6x10^7^ cells/mL) in the center of 48-well plate and cultured under chondrogenic condition for 3 days. (Bars show mean ± SEM. n=6∼9, *p<0.05, **p<0.01, *** p<0.005, **** p<0.0001).

## Discussion

Being the most commonly prescribed diabetic medication in the world, metformin’s effects on bone healing in T2D patients remain unclear. To assess the impacts of metformin on fracture healing under hyperglycemic condition, we applied several classic bone fracture models in T2D mice. Our results demonstrated that in all injury models tested, metformin successfully rescued the delayed bone healing and remolding in T2D mice by facilitating bone formations. Further cell culture studies demonstrated the mechanism of metformin’s action at cellular level via promoting the proliferation, differentiation, and lineage commitment of primary BMSCs. Taken together, metformin showed its potential as an effective drug for increasing the rate and success of bone healing in diabetic patients that are not taking metformin on regular basis.

In all the bone fracture models studied in this study, metformin significantly enhanced bone-healing parameters in MKR mice. However, in the WT animals, quantitative analysis of images showed no difference between the PBS and the metformin treatment in terms of bone healing. These data suggest that metformin is only beneficial for bone healing under hyperglycemic conditions but does not enhance bone healing in WT animals without diabetes. In all BMSC based essays, metformin was not administered to culture media in vitro but administered to animals in vivo before the BMSC were isolated. Administration of metformin in MKR mice successfully salvaged the BMSC proliferation capability and lineage commitment and brought it back to the equivalent level as observed in WT group. Interestingly, metformin did not affect the proliferation and lineage commitment of BMSCs in WT group when compared to the PBS vehicle. These data are consistent with those obtained from bone fracture models, suggesting that metformin may exert its effects through normalizing hyperglycemia (Fig. S1A), glucose tolerance (Fig. S1B) and other metabolic disturbance under diabetic conditions and does not enhance bone healing in WT animals. In ovariectomized rats, impaired bone density and quality were significantly improved by the treatment of metformin [6]. Taken together, it seems that metformin does not promote further bone growth under physiological conditions but helps to maintain bone homeostasis under pathological conditions such as hyperglycemia and lack of estrogen. To our knowledge, there are very scarce amount of research on the direct effects of metformin on BMSCs. In an in vitro study using bone marrow-derived multipotent mesenchymal stromal cells, metformin causes inhibition of proliferation and abnormalities of their morphology and ultrastructure [18]. This might be due the level of metformin used in the culture system or that metformin does not work directly on BMSCs but requires other tissues, cells and the in vivo context to exert its effect in mice. In another paper, metformin added to culture is shown to promote the osteogenesis of BMSCs isolated from T2D patients and osseointegration when administered in rats[19].Whether metformin has direct beneficial effects on BMSCs remains to be investigated.

As reviewed by Roszer, T. [20], diabetes is accompanied by increased level of pro-inflammatory factors, reactive oxygen species (ROS) generation and accumulation of advanced glycation end products (AGEs). The increased inflammatory state could result in apoptosis of osteoblasts and prolonged survival of osteoclasts, which lead to early destruction of callus tissue and impair bone fracture healing of diabetic patients. Thus, antagonizing inflammatory signal pathways and inhibition of inflammation may deserve greater attention in the management of diabetic fracture healing. There are substantial evidence supporting that metformin not only improves chronic inflammation by attenuating hyperglycemia but also has a direct anti-inflammatory effect. Targeting inflammatory pathways seems to be an important part of the comprehensive mechanisms of action of this drug [21]. In addition to AMPK activation and inhibition of mTOR pathways, metformin acts on mitochondrial function and cellular homeostasis processes such as autophagy [21]. Both dysregulated mitochondria and failure of the autophagy pathways affect cellular health drastically and can trigger the onset of metabolic and age-associated inflammation and diseases. For example, T-helper type 17 (Th17) cells, an important proinflammatory CD4+ T cell subset secreting interleukin 17 (IL-17), has been suggested to play an essential role in development of diabetes mellitus [22]. Metformin can ameliorate the pro-inflammatory profile of Th17 by increasing autophagy and improving mitochondrial bioenergetics [23]. In addition, at day 21 post fracture, the PGC1α in callus tissues isolated from the fracture site was significantly upregulated in metformin treated MKR mice (Fig. S2H). As reviewed by Halling and Pilegaard [24], PGC-1α not only regulates mitochondrial biogenesis but also its function. PGC-1α- mediated regulation of mitochondrial quality may contribute to many age-related dysfunctions including insulin sensitivity. Anti-inflammation and enhancement of mitochondrial function could be very important means that metformin utilizes to facilitate bone formation and healing under hyperglycemic conditions.

Diabetic hyperglycemia has been suggested to play a role in osteoarthritis. The metabolic alterations in body fluid such as hyperglycemia could negatively affect the cartilage through direct effects on chondrocytes by stimulating the production of advanced glycosylation end products (AGEs) accumulation in the synovium [25]. PPARγ is highly expressed in adipocytes and the downregulation of PPARγ expression in the callus of metformin treated MKR mice reflected the shift of mesenchymal cells fate. In T2DM mouse model, differentiation of growth plate chondrocytes is delayed and this delay may result from premature apoptosis of the growth plate chondrocytes [26]. Besides its effects on bone formation, there are also interests in studying the effects of metformin on chondrocytes, especially in the context of osteoarthritis development. Limited reference showing that metformin is protective against development of osteoarthritis by reducing chondrocyte apoptosis and alleviating chondrocyte degeneration [27-29]. Consistent with the above reports, our data suggest that metformin promoted the cartilage formation in the endochondral ossification at day 14 post-fracture in T2D mice. Moreover, metformin rescued the chondrocyte disc formation in BMSCs isolated from T2D mice when compared to the PBS treated control. Metformin also upregulated chondrocyte transcript factor SOX9 and in callus tissue isolated at the fracture site in metformin treated MKR mice. Sox-9 plays an essential role in regulation of cartilage matrix production and cartilage repair [15]. In addition, PGC1α was significantly upregulated in metformin treated MKR mice when compared to the PBS-treated MKR animals.

PGC1alpha is important to maintain chondrocyte metabolic flexibility and tissue homeostasis. The loss of PGC1α in chondrocytes during OA pathogenesis resulted in the activation of mitophagy and stimulated cartilage degradation and apoptotic death of chondrocytes [30]. The activation of PGC1α is a potential strategy to delay or prevent the development of OA. The cellular signaling pathways through which metformin exerts its protective effects in chondrocytes warrant further research.

In conclusion, our study demonstrated that metformin can facilitate bone healing, bone formation and chondrogenesis in T2D mice. The molecular mechanism of metformin’s action demands further research in hope to identify specific therapeutic target to facilitate bone healing and repair in diabetic patients.

## Conflict of interest disclosure

Authors declare no conflict of interest.

## ACKNOWLEDGMENTS

This study was funded by New York University Start-up and Career Enhancement Award to XL. The authors would like to thank the support from the microCT core facility at New York University College of Dentistry.

## Supplementary Materials

### Methods

#### Glucose test and Glucose tolerance test

WT and MKR mice of 12-week-old were treated with PBS or metformin through daily intraperitoneal injection as previously described. Blood glucose level was recorded at indicated time points for each mice.

After 14 days PBS or metformin injection, animals were fasting overnight for 16 hours by transferring into a clean new cage supplied with water. 20% (W/V) glucose solution was applied to animals through oral gavage (1.6g of glucose/ KG body weight). Blood glucose was measured at indicated time points till two hours post glucose oral gavage.

#### Quantitative PCR

Femoral closed fracture was performed as previously described. Callus tissue at the fracture cite was collected and lysate in Trizol Reagent. RNA was extracted and purified using the RNeasy Plus kit. cDNA was prepared by using a Reverse Transcriptase Kit, and PCR was then performed with a Bio-Rad CFX384 Real-Time PCR detection System.

Primers information as below: mβ-actin Fr: AGCCATGTACGTAGCCATCC; mβ-actin Rv: CTCTCAGCTGTGGTGGTGAA; mPGC1α Fr: GAATCAAGCCACTACAGACACCG; mPGC1α Rv: CATCCCTCTTGAGCCTTTCGTG; mSP7Fr: GGCTTTTCTGCGGCAAGAGGTT; mSP7 Rv: CGCTGATGTTTGCTCAAGTGGTC; mRUNX2 Fr: CCTGAACTCTGCACCAAGTCCT; mRUNX2 Rv: TCATCTGGCTCAGATAGGAGGG; mPPARγ Fr: GTACTGTCGGTTTCAGAAGTGCC; mPPARγ Rv: ATCTCCGCCAACAGCTTCTCCT; mC/EBPα Fr: CAAAGCCAAGAAGTCGGTGGACAA; mC/EBPα Rv: TCATTGTGACTGGTCAACTCCAGC; mSOX9 Fr: AAGAAAGACCACCCCGATTACA; mSOX9 Rv: CAGCGCCTTGAAGATAGCATT.

---

**Figure S1.**
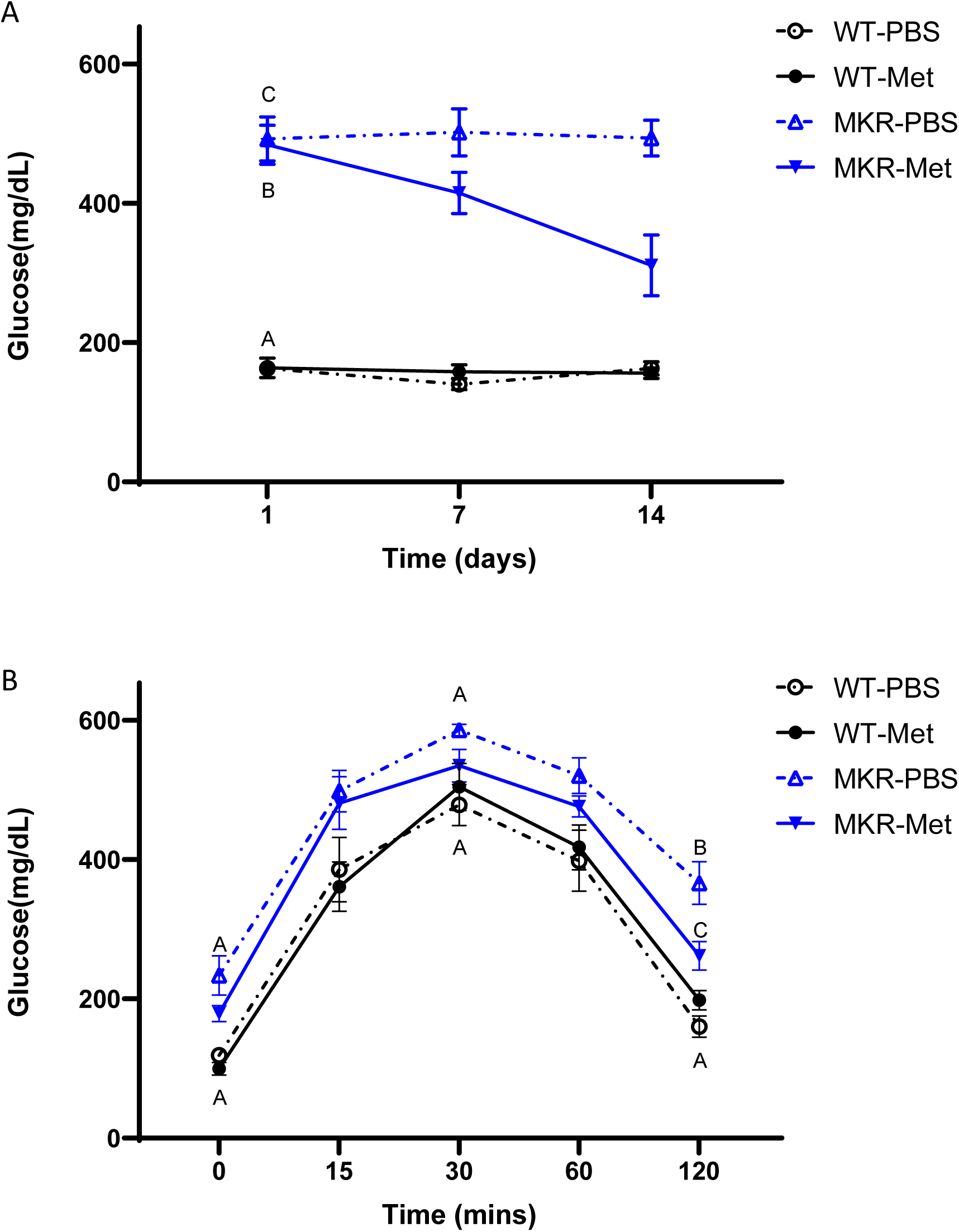
Metformin’s effect on glucose levels in MKR mice. A) Glucose levels in WT and MKR mice throughout the 14 days PBS or metformin treatment. B) Glucose tolerance test (GTT) after fasting post 14 days PBS or metformin treatment. (N=6, SD) If different letters are shown at a time point, they are statistically different from one another (ANOVA, p<0.05 by post-hoc Tukey’s).

**Figure S2.**
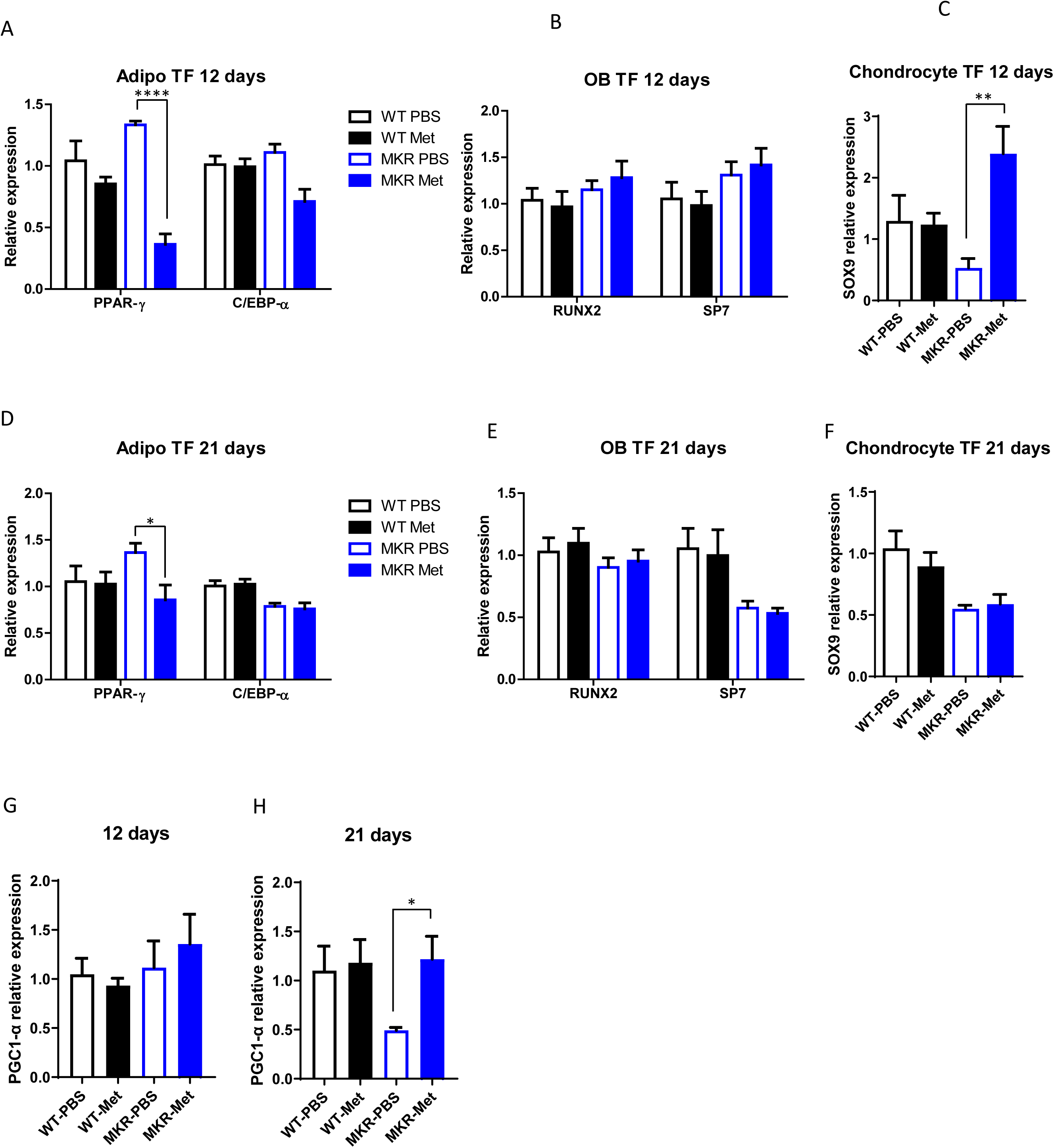
Adipocyte and chondrocyte transcript factors expression within the femoral fracture callus tissue. A) adipocyte, B) osteoblast differentiation, and C) chodrogenesis transcript factors expression in the callus tissue at 12 days post-fracture. D) adipocyte, E) osteoblast differentiation, and F) chodrogenesis transcript factors expression in the callus tissue at 21 days post-fracture. PGC1α expression in the callus tissue at G) 12 days H) 21 days post-fracture.

## Notes

### Competing Interest Statement

The authors have declared no competing interest.

## References

[1] G. Danaei, M.M. Finucane, Y. Lu, G.M. Singh, M.J. Cowan, C.J. Paciorek, J.K. Lin, F. Farzadfar, Y.H. Khang, G.A. Stevens, M. Rao, M.K. Ali, L.M. Riley, C.A. Robinson, M. Ezzati, G. Global Burden of Metabolic Risk Factors of Chronic Diseases Collaborating, National, regional, and global trends in fasting plasma glucose and diabetes prevalence since 1980: systematic analysis of health examination surveys and epidemiological studies with 370 country-years and 2.7 million participants, Lancet, 378 (2011) 31-40.

[2] W. Yan, X. Li, Impact of diabetes and its treatments on skeletal diseases, Frontiers of medicine, 7 (2013) 81–90.

[3] P. Vestergaard, L. Rejnmark, L. Mosekilde, Relative fracture risk in patients with diabetes mellitus, and the impact of insulin and oral antidiabetic medication on relative fracture risk, Diabetologia, 48 (2005) 1292–1299.

[4] Y. Gao, J. Xue, X. Li, Y. Jia, J. Hu, Metformin regulates osteoblast and adipocyte differentiation of rat mesenchymal stem cells, J Pharm Pharmacol, 60 (2008) 1695–1700.

[5] M. Shah, B. Kola, A. Bataveljic, T.R. Arnett, B. Viollet, L. Saxon, M. Korbonits, C. Chenu, AMP-activated protein kinase (AMPK) activation regulates in vitro bone formation and bone mass, Bone, 47 (2010) 309–319.

[6] Y. Gao, Y. Li, J. Xue, Y. Jia, J. Hu, Effect of the anti-diabetic drug metformin on bone mass in ovariectomized rats, Eur J Pharmacol, 635 (2010) 231–236.

[7] M.S. Molinuevo, L. Schurman, A.D. McCarthy, A.M. Cortizo, M.J. Tolosa, M.V. Gangoiti, V. Arnol, C. Sedlinsky, Effect of metformin on bone marrow progenitor cell differentiation: in vivo and in vitro studies, J Bone Miner Res, 25 (2010) 211–221.

[8] F.T.T. Lai, B.H.K. Yip, D.J. Hunter, D.P. Rabago, C.D. Mallen, E.K. Yeoh, S.Y.S. Wong, R.W. Sit, Metformin use and the risk of total knee replacement among diabetic patients: a propensity-score-matched retrospective cohort study, Sci Rep, 12 (2022) 11571.

[9] A.M. Fernandez, J.K. Kim, S. Yakar, J. Dupont, C. Hernandez-Sanchez, A.L. Castle, J. Filmore, G.I. Shulman, D. Le Roith, Functional inactivation of the IGF-I and insulin receptors in skeletal muscle causes type 2 diabetes, Genes & development, 15 (2001) 1926–1934.

[10] L.C. Gerstenfeld, T.J. Cho, T. Kon, T. Aizawa, A. Tsay, J. Fitch, G.L. Barnes, D.T. Graves, T.A. Einhorn, Impaired fracture healing in the absence of TNF-alpha signaling: The role of TNF-alpha in endochondral cartilage resorption, Journal of Bone and Mineral Research, 18 (2003) 1584–1592.

[11] F. Bonnarens, T.A. Einhorn, Production of a standard closed fracture in laboratory animal bone, J Orthop Res, 2 (1984) 97–101.

[12] R.M. Voigt, C.B. Forsyth, S.J. Green, P.A. Engen, A. Keshavarzian, Circadian Rhythm and the Gut Microbiome, Int Rev Neurobiol, 131 (2016) 193–205.

[13] T.A. Einhorn, L.C. Gerstenfeld, Fracture healing: mechanisms and interventions, Nat Rev Rheumatol, 11 (2015) 45–54.

[14] C.J. Rosen, The epidemiology and pathogenesis of osteoporosis, Endotext [Internet], (2020).

[15] E. Kolettas, H.I. Muir, J.C. Barrett, T.E. Hardingham, Chondrocyte phenotype and cell survival are regulated by culture conditions and by specific cytokines through the expression of Sox-9 transcription factor, Rheumatology (Oxford), 40 (2001) 1146–1156.

[16] Y. Kawakami, M. Tsuda, S. Takahashi, N. Taniguchi, C.R. Esteban, M. Zemmyo, T. Furumatsu, M. Lotz, J.C. Izpisua Belmonte, H. Asahara, Transcriptional coactivator PGC-1alpha regulates chondrogenesis via association with Sox9, Proc Natl Acad Sci U S A, 102 (2005) 2414–2419.

[17] X. Zhao, F. Petursson, B. Viollet, M. Lotz, R. Terkeltaub, R. Liu-Bryan, Peroxisome proliferator-activated receptor gamma coactivator 1alpha and FoxO3A mediate chondroprotection by AMP-activated protein kinase, Arthritis Rheumatol, 66 (2014) 3073–3082.

[18] A. Smieszek, A. Czyrek, K. Basinska, J. Trynda, A. Skaradzinska, A. Siudzinska, M. Maredziak, K. Marycz, Effect of Metformin on Viability, Morphology, and Ultrastructure of Mouse Bone Marrow-Derived Multipotent Mesenchymal Stromal Cells and Balb/3T3 Embryonic Fibroblast Cell Line, Biomed Res Int, 2015 (2015) 769402.

[19] R. Sun, C. Liang, Y. Sun, Y. Xu, W. Geng, J. Li, Effects of metformin on the osteogenesis of alveolar BMSCs from diabetic patients and implant osseointegration in rats, Oral Dis, 28 (2022) 1170–1180.

[20] T. Roszer, Inflammation as death or life signal in diabetic fracture healing, Inflamm Res, 60 (2011) 3–10.

[21] L.P. Bharath, B.S. Nikolajczyk, The intersection of metformin and inflammation, Am J Physiol Cell Physiol, 320 (2021) C873–C879.

[22] A. Abdel-Moneim, H.H. Bakery, G. Allam, The potential pathogenic role of IL- 17/Th17 cells in both type 1 and type 2 diabetes mellitus, Biomed Pharmacother, 101 (2018) 287–292.

[23] L.P. Bharath, M. Agrawal, G. McCambridge, D.A. Nicholas, H. Hasturk, J. Liu, K. Jiang, R. Liu, Z. Guo, J. Deeney, C.M. Apovian, J. Snyder-Cappione, G.S. Hawk, R.M. Fleeman, R.M.F. Pihl, K. Thompson, A.C. Belkina, L. Cui, E.A. Proctor, P.A. Kern, B.S. Nikolajczyk, Metformin Enhances Autophagy and Normalizes Mitochondrial Function to Alleviate Aging-Associated Inflammation, Cell Metab, 32 (2020) 44–55 e46.

[24] J.F. Halling, H. Pilegaard, PGC-1alpha-mediated regulation of mitochondrial function and physiological implications, Appl Physiol Nutr Metab, 45 (2020) 927–936.

[25] Q. Li, Y. Wen, L. Wang, B. Chen, J. Chen, H. Wang, L. Chen, Author Correction: Hyperglycemia-induced accumulation of advanced glycosylation end products in fibroblast-like synoviocytes promotes knee osteoarthritis, Exp Mol Med, 54 (2022) 862–865.

[26] K. Wongdee, N. Charoenphandhu, Update on type 2 diabetes-related osteoporosis, World J Diabetes, 6 (2015) 673–678.

[27] J. Yan, G. Feng, L. Ma, Z. Chen, Q. Jin, Metformin alleviates osteoarthritis in mice by inhibiting chondrocyte ferroptosis and improving subchondral osteosclerosis and angiogenesis, J Orthop Surg Res, 17 (2022) 333.

[28] J. Yan, D. Ding, G. Feng, Y. Yang, Y. Zhou, L. Ma, H. Guo, Z. Lu, Q. Jin, Metformin reduces chondrocyte pyroptosis in an osteoarthritis mouse model by inhibiting NLRP3 inflammasome activation, Exp Ther Med, 23 (2022) 222.

[29] X. Feng, J. Pan, J. Li, C. Zeng, W. Qi, Y. Shao, X. Liu, L. Liu, G. Xiao, H. Zhang, X. Bai, D. Cai, Metformin attenuates cartilage degeneration in an experimental osteoarthritis model by regulating AMPK/mTOR, Aging (Albany NY), 12 (2020) 1087–1103.

[30] D. Kim, J. Song, E.J. Jin, BNIP3-Dependent Mitophagy via PGC1alpha Promotes Cartilage Degradation, Cells, 10 (2021).

